# Tryptophan-based motifs in the LLP3 Region of the HIV-1 envelope glycoprotein cytoplasmic tail direct trafficking to the endosomal recycling compartment and mediate particle incorporation

**DOI:** 10.1101/2023.04.28.538708

**Authors:** Grigoriy Lerner, Lingmei Ding, Paul Spearman

## Abstract

The HIV-1 envelope glycoprotein complex (Env) is incorporated into developing particles at the plasma membrane (PM). The cytoplasmic tail (CT) of Env is known to play an essential role in particle incorporation, while the exact mechanisms underlying this function of the CT remain uncertain. Upon reaching the PM, trafficking signals in the CT interact with host cell endocytic machinery, directing Env into endosomal compartments within the cell. Prior studies have suggested that Env must traffic through the endosomal recycling compartment (ERC) in order for Env to return to the plasma membrane (PM) site of particle assembly. Expression of a truncated form of the ERC-resident trafficking adaptor Rab11-Family Interacting Proteins C (FIP1C) resulted in CT-dependent sequestration of Env in the condensed ERC, preventing recycling of Env to the PM. In this work, the motifs within the CT responsible for ERC localization of Env were systematically mapped. A small deletion encompassing the N-terminal portion of LLP3 eliminated ERC localization. Site-directed mutagenesis identified two tryptophan-based motifs (WE_790-791_ and WW_796-797_) within the N-terminus of LLP3 that were essential for ERC localization of Env. Mutant viruses bearing substitutions in these motifs were deficient in Env incorporation, with a corresponding loss of particle infectivity and a significant defect in replication in a spreading infection assay. These results identify two tryptophan-based motifs at the N-terminal portion of LLP3 that mediate ERC localization and Env incorporation, providing additional supporting evidence for the importance of cellular recycling pathways in HIV-1 particle assembly.

**IMPORTANCE:** The HIV-1 envelope glycoprotein (Env) is an essential component of the virus, and has an exceedingly long cytoplasmic tail (CT). Previous studies have suggested that trafficking signals in the CT interact with host factors to regulate the incorporation of Env into particles. One particular area of interest is termed lentiviral lytic peptide 3 (LLP3), as small deletions in this region have been shown to disrupt Env incorporation. In this study, we identify a small region within LLP3 that regulates how Env associates with cellular recycling compartments. Mutants that reduced or eliminated Env from the recycling compartment also reduced Env incorporation into particles. These findings emphasize the importance of two tryptophan motifs in LLP3 to the incorporation of Env into particles, and provide additional support for the idea that the CT interacts with host recycling pathways to determine particle incorporation.

## INTRODUCTION

The envelope glycoprotein (Env) of HIV-1 is an essential viral protein that mediates binding to CD4 and coreceptor molecules, thereby triggering fusion and entry into target cells. Env is also the principal neutralizing determinant of the virus and has been the focus of active research in the HIV vaccine field. Env is synthesized on the rough ER as the precursor protein gp160, where it is co-translationally glycosylated and assembled into trimers (1, 2). Env is cleaved by furin-like proteases in the Golgi apparatus into the gp120 surface (SU) and gp41 transmembrane subunits (TM), which remain non-covalently associated as the heterotrimer is transported to the plasma membrane (3, 4). Upon arrival at the plasma membrane, Env trimers are rapidly endocytosed via a clathrin-and AP-2-dependent mechanism that relies upon a membrane-proximal Yxx< motif in the Env CT (5–9). The trafficking steps taken by Env following endocytosis that regulate its anterograde movement back to the PM for particle incorporation remain incompletely defined.

Host recycling pathways are likely to be required for directing Env to the site of particle assembly and promoting its incorporation into developing HIV-1 particles. FIP1C is a trafficking adaptor that has been implicated in trafficking of Env from the ERC to the plasma membrane. Depletion of FIP1C led to a reduction of Env incorporation in T cell lines, and overexpression of a C-terminal fragment of FIP1C resulted in sequestration of Env within the ERC (10, 11). The small GTPase Rab14 interacts with FIP1C and contributes to Env recycling and particle incorporation (11, 12). An alternative pathway for endocytosed Env involves interaction with the retromer complex, resulting in retrograde trafficking of Env back to the Golgi and leading to a reduction of Env at the cell surface (13). A fraction of intracellular Env has also been observed to colocalize with markers of the lysosome (6, 14, 15). Inhibitors of lysosomal function result in reduced degradation of Env, making it likely that lysosomal degradation is the default fate of endocytosed Env in the absence of active anterograde trafficking (3, 15). Thus, multiple potential trafficking pathways are available for Env trafficking following endocytosis, with some routes leading to productive particle incorporation at the PM, and others directing Env to alternative pathways including degradation in the lysosome. Identification of specific trafficking motifs present in the CT that are responsible for recycling and subsequent particle incorporation are therefore of considerable interest, and are the subject of this report.

The presence of a long CT is a characteristic feature of most lentiviral Env proteins. HIV-1 and SIV incorporate a CT of approximately 150 amino acids. The expression of a long CT suggests that this provides an added function or functions that are evolutionarily advantageous to the virus (16). The Env CT consists of an N-terminal unstructured region followed by three amphipathic alpha helices known as the lentiviral lytic peptides (LLPs), numbered from N-to C-terminus as LLP2/LLP3/LLP1 (17). These domains have been shown to contribute to a baseplate structure supported by a complex network of intratrimer bonds, as well as bonds linking the LLPs to the transmembrane domain (18, 19). Specific motifs within the LLPs have previously been implicated in interactions with cellular trafficking machinery. Interactions with the retromer complex have been mapped to regions of the CT termed inhibitory sequences (IS1 and IS2), with IS1 in the unstructured region of the CT and IS2 overlapping LLP2 (13, 20). The N-terminal portion of LLP3 appears to have particular importance for Env incorporation into HIV-1 particles. Murakami and Freed showed that small deletions in this region result in Env incorporation defects and reduce viral replication (21). A systematic mutagenesis of tyrosine-based and dileucine motifs in the CT found that a nine-residue sequence at the N-terminus of LLP3 (Y_795_WWNLLQYW_802_) plays a critical role in HIV replication in T cells (22). Our group previously demonstrated that the YW_795_ motif is an important determinant of Env incorporation, as a YW_795_SL mutant virus was deficient in Env incorporation and replicated poorly in T cell lines (23, 24). The Env incorporation defect introduced by mutation of the YW_795_ motif could be rescued by a downstream second-site reversion introducing a single amino acid change near the C-terminus of the CT (YW_795_SL/L_850_S), restoring Env incorporation and viral replication. Although a direct interaction of this motif with FIP1C has not been demonstrated, the YW_795_SL mutant virus was insensitive to FIP1C depletion, while Env incorporation by revertant YW_795_SL/L_850_S was reduced by FIP1C depletion. Other groups have examined the second YW motif in this region, YW_802_, showing that disruption of this motif reduces Env incorporation and fusogenicity (22, 25, 26).

An intriguing aspect of Env incorporation into particles is that the CT is required for particle incorporation in a cell type-dependent manner (27). Large truncations or deletions of the CT do not inhibit the incorporation of Env into particles budding from model epithelial cell lines including 293T and COS cells, or from the MT-4 T cell line. Cell types that do not require an intact CT are termed permissive cells, indicating the permissive nature of incorporation of Env bearing a truncated CT. In contrast, an intact CT is required for efficient particle incorporation and replication in many T cell lines including Jurkat, CEM, and H9 (27–29). Furthermore, an intact CT is required for particle incorporation and replication in primary CD4+ T cells and macrophages (27). The precise mechanism by which the CT mediates HIV-1 Env incorporation remains incompletely understood. However, a growing body of evidence supports a role for intracellular trafficking mediated by the CT in directing Env incorporation into HIV-1 particles (reviewed in (30)). Notably, the YW_795_SL mutant in LLP3 reproduced the phenotype of CT144, with efficient incorporation in permissive cells and poor incorporation in nonpermissive cells (24). This suggests that trafficking motifs in the N-terminal portion of LLP3 may be responsible for the cell type-dependent incorporation of Env.

We previously reported that expression of a C-terminal fragment of FIP1C (FIP1C_560-649_) retained or “trapped” Env in a perinuclear compartment, with a corresponding reduction in the incorporation of Env into particles (10). Trapping of Env by FIP1C_560-649_ was dependent on the presence of the CT, and resulted in prominent Env colocalization with ERC markers such as Rab11a and Rab14 (10). This suggested to us that FIP1C_560-649_ colocalization could be used to map motifs in the CT that mediate ERC interactions and subsequent recycling events. In the present study, we define specific residues within the Env CT that are required for trafficking to/retention in the ERC. Using CT truncation followed by site-directed mutagenesis, we found that residues adjacent to the LLP2-LLP3 junction are required for ERC retention of Env. Mutation of two tryptophan-based motifs in the N-terminal portion of LLP3 (WE_790-791_ and WW_796-797_) resulted in a significant loss of particle incorporation, and viruses bearing substitutions in these motifs replicated very poorly in T cell lines. Thus, the tryptophan motifs in the N-terminal portion of LLP3 mediate both ERC localization and particle incorporation, emphasizing the importance of host recycling pathways in determining particle incorporation of Env.

## RESULTS

### Mapping Determinants in CT required for ERC localization of Env

To define which region of the CT plays a role in ERC localization, we initially constructed a series of Env proteins in which the structural Lentiviral Lytic Peptides (LLPs) were deleted from C-to N-terminus, beginning with the most C-terminal (Δ1CT33), then Δ1CT71, and finally Δ1CT104 (Figure 1A, with position of stop codon shown in 1B). An additional construct, Δ1CT82, extended the deletion into LLP2 with a stop codon following residue 774 (relative to HXB2 reference strain). We then coexpressed each of the truncated Env proteins shown in Figure 1 with FIP1C_560-649_, and assessed colocalization of Env with FIP1C_560-649_ as a measure of ERC localization. As described previously (10), WT Env strongly colocalized with FIP1C_560-649_ in the perinuclear ERC (Figure 2A, top panel), while Env bearing a truncation of 144 residues of the CT did not (Figure 2A, Δ1CT144). Deletion of the C-terminal LLP1 alpha helix resulted in more Env localized outside of the ERC than WT, along with a partial reduction in colocalization (Figure 2A, Δ1CT33, with quantitation in Figure 2B). CT truncation that resulted in the loss of two C-terminal LLPs (LLP3 and LLP1) resulted in a more severe loss of colocalization, suggesting an important role for determinants in LLP3 (Figure 2A and 2B, Δ1CT71). Complete loss of colocalization was observed when the truncation was performed within the mid-portion of LLP2, resulting in loss of the 82 residues following L_774_ (Figure 2A and 2B, Δ1CT82). A truncation that resulted in loss of all three LLPs also showed complete loss of colocalization, similar to results seen with Δ1CT82 and with Δ1CT144 (Figure 2A, and 2B, Δ1CT104). The sequential loss of ERC colocalization is depicted in Figure 2B, illustrating that the loss of the C-terminal 82 residues of the CT was sufficient to completely prevent ERC localization. These CT truncation results support a model in which the LLPs play a role in Env trafficking to the ERC. The LLP2/LLP3 junction was selected for further studies, as complete loss of colocalization required both LLP3/LLP1 deletion and the C-terminal portion of LLP2 represented by ΔCT82.

**FIG 1.**
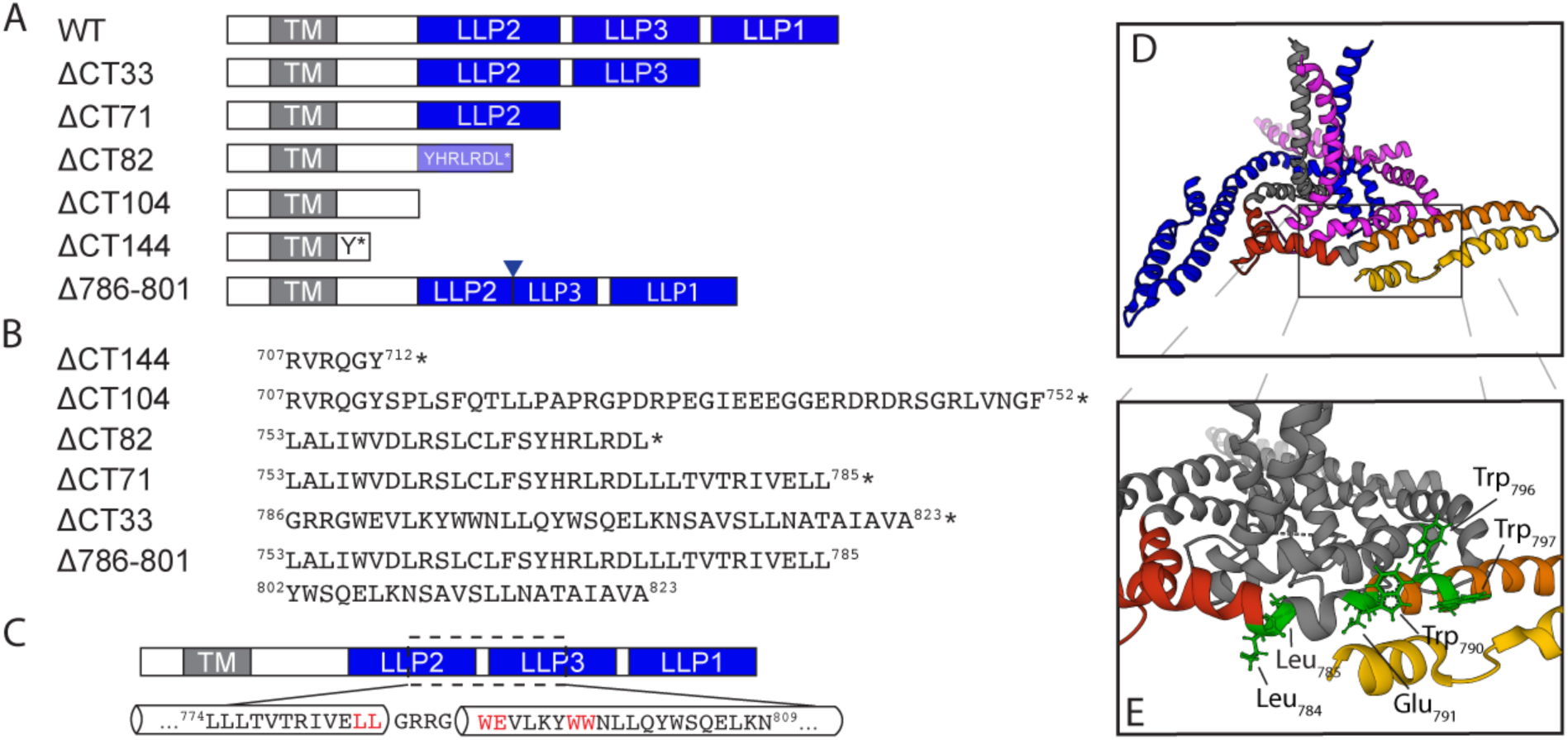
Env mutants used for this study. (A) Schematic representation of the truncation mutants used in this study with respect to structural elements within the CT. The mutants are named for the number of amino acids removed the C-terminus of the CT. (B) The precise location of stop codons in the sequence of each truncation mutant is shown with the preceding LLP or other structural element, asterisk indicates position of stop codon. (C) Depiction of LLP2-3 region which was selected for alanine scanning mutagenesis. Amino acids highlighted in red were found to be important for ERC localization of Env. (D) Structure of the HIV-1 Env transmembrane region coupled to the baseplate as proposed by Piai et al., image created using PyMol software using Mol* on the RCSB PDB site and PDB file 7LOH (19). The three trimer subunits are depicted in blue, gray, and magenta, while the CT of the gray subunit is further subdivided red, orange, and yellow for LLP2, LLP3, and LLP1 respectively. (E) location of amino acid residues highlighted in (C) mapped onto the structure in (D).

**FIG 2.**
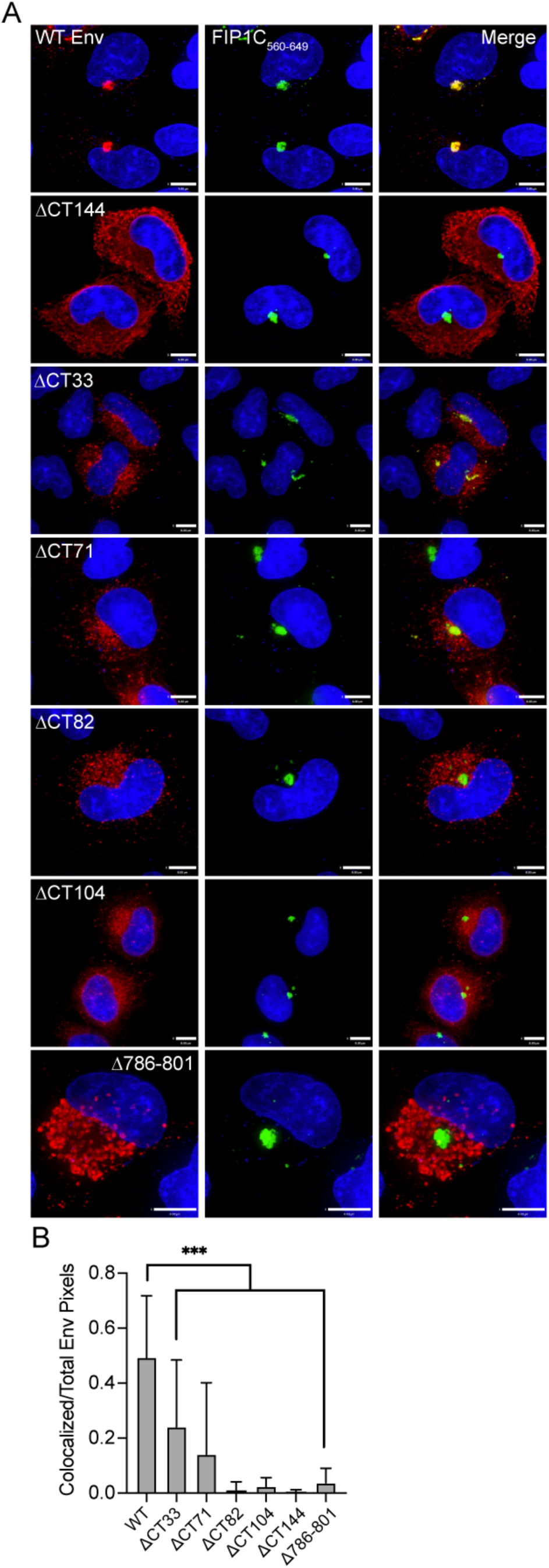
Colocalization of Envelope with FIP1C560-649 requires LLP3 and the C-terminal segment of LLP2. (A) HeLa cells were transfected with WT or truncated Env and GFP-FIP1C_560-649_ for 48 hours. Cells were then fixed, permeabilized, and stained with human anti-gp120 antibody 2G12, shown in red. GFP-FIP1C_560-649_ representing the condensed ERC is shown in green. Bar = 8 μm. (B) Colocalization of WT or truncated Env with GFP-FIP1C_560-649_ was quantified from 20 cells per Env mutant across more than three separate experiments; Manders colocalization coefficient is reported as mean ± SD. Significance was assessed using One-way ANOVA with Dunnett correction for multiple comparisons. *** P<0.001.

In order to further evaluate the importance of this region in mediating ERC localization of Env, we next created an internal deletion of residues 786-801, representing the very N-terminal segment of LLP3 (Figure 1B, colored text) while keeping LLP2 and LLP1 intact. Remarkably, this mutant was completely excluded from the ERC (Figure 2A, with quantitation shown in 2B). We noted a subtle difference in subcellular distribution of 1′786-801 from that of ΔCT144, with Δ786-801 present in larger compartments outside of the ERC, consistent with defective Env sorting and failure to reach the ERC. The fact that this internal deletion at the N-terminus of LLP3 disrupted ERC localization and perturbed Env distribution reinforces the importance of this region in intracellular trafficking of Env.

### Cell type-specific differences in incorporation of Env with disruption of LLP3

We next examined incorporation of truncated Env into released particles, testing the hypothesis that mutants that were unable to reach the ERC would be deficient in particle incorporation. To do so, we generated truncation mutants within the NL4-3 proviral expression plasmid by inserting stop codons at the specific locations shown in Figure 1B. All Env deletion constructs in the proviral context resulted in release of virus from transfected 293T or HeLa cells at a level comparable to WT as measured by p24 ELISA (data not shown). Env incorporation was assessed by Western blotting of viruses produced from transfected HeLa cells and from H9 cells infected with VSV-G-pseudotyped viruses. Note that we chose these two cell types in order to contrast a semipermissive cell type (HeLa) with that of a nonpermissive cell type (H9). As expected, 1′CT144 Env was incorporated into particles produced from transfected HeLa cells, although to a lesser extent than WT (Figure 3A and 3C). Deletion of all three LLPs (1′CT104, Figure 3A) allowed Env incorporation at levels slightly above that of 1′CT144 (Figure 3C). Truncation of the CT prior to the C-terminal LLP1 domain (ΔCT33) reduced Env incorporation to levels similar to that of CT144. Surprisingly, deletion of LLP1 and LLP3 while maintaining all (ΔCT71) or just the N-terminal portion of LLP2 (ΔCT82) completely eliminated Env incorporation into particles produced from HeLa cells, without disrupting cellular levels of Env (Figure 3A and 3C). These results, showing a loss of Env incorporation for truncations in LLP2 and LLP3 but not for more drastic truncations (1′CT144, ΔCT104) or less drastic mutations (ΔCT33) are reminiscent of findings previously observed by the Aiken lab (31). In contrast to HeLa cells, when these same CT truncated viruses were produced from infected H9 cells, all truncations resulted in near complete defects in Env incorporation (Figure 3B). This suggests that cell type-specific differences in Env incorporation exist that can be mapped within the CT, and that the LLP2/3 junction may be key to understanding the differences seen in nonpermissive vs. permissive cells.

**FIG 3.**
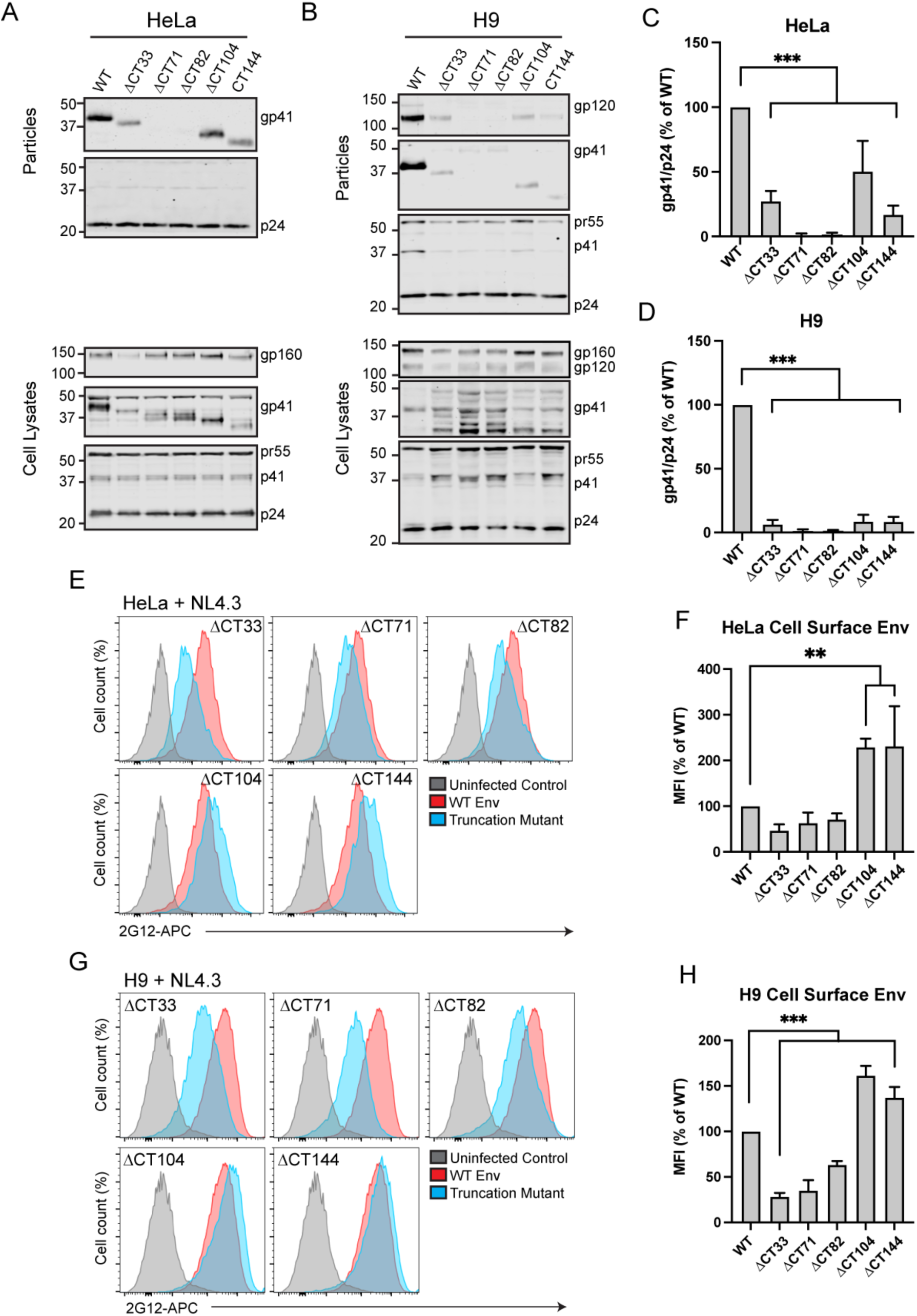
Truncation of Env restricts incorporation in both permissive and restrictive cell types. (A, B) Viral supernatants were produced from transfected HeLa cells or H9 cells infected with VSV-G-pseudotyped NL4-3 virus encoding truncated Env. Supernatants were pelleted through a sucrose cushion, normalized for virus production by p24 ELISA, and subjected to immunoblotting. (C, D) Westerns were analyzed by densitometry and gp41/p24 ratios from four experiments are reported as percent of WT. (E) HeLa cells transfected with NL4-3 truncation mutants were analyzed for Env levels on the cell surface by staining of fixed, unpermeabilized cells with 2G12 directly conjugated to APC. Cells were then permeabilized and stained for Gag to gate for infected cells. (F) MFI values were normalized to WT and averaged for three experiments. Results are reported as mean ± SD. G) H9 cells infected with VSV-G-pseudotyped NL4-3 mutants were assessed for cell surface Env as described in E. (H) MFI values from H9 cells represented as described for F. Significance was assessed for truncation mutants vs WT control using one-way ANOVA with Dunnett correction for multiple comparisons. **, P<0.01; ***, P<0.001.

The cell type-specific Env incorporation differences with CT truncation mutants described above were not completely explained by levels of cell surface Env. In HeLa cells, removal of LLP1 (ΔCT33) resulted in a reduction of cell surface Env down to 47% of WT, while ΔCT71 reduced cell surface Env to 63% of WT and ΔCT82 dropped Env to 71% of WT (Figure 3E and 3F). In contrast, ΔCT104 and ΔCT144 were significantly enriched on the cell surface relative to WT, with a 229% increase in MFI for ΔCT104 and 231% increase in MFI for ΔCT144 (Figure 3E and 3F). Findings in H9 cells were similar to HeLa, with diminished cell surface Env for ΔCT33, ΔCT71, and ΔCT82 (28%, 35%, and 63% of WT, respectively) and increased cell surface Env for ΔCT104 and CT144 (161% and 137%, respectively, Figure 3G and 3H). Thus, truncations removing LLP1, LLP1 and LLP3, or part of LLP2 in conjunction with LLP3 and LLP1 reduced cell surface Env, while complete removal of all three LLPs or truncation of the C-terminal 144 residues led to significantly increased cell surface envelope levels in both cell types. One interpretation of these findings is that removal of CT motifs relevant to Env trafficking and particle incorporation reduces incorporation of Env in nonpermissive cells, as reflected in poor Env incorporation even in the presence of enhanced cell surface Env in the H9 T cell line. In Hela cells, however, significant increases in cell surface levels resulting from LLP2/LLP3/LLP1 deletion or for ΔCT144 are able to overcome this trafficking defect through another mechanism, such as through passive incorporation when there are increased levels of Env at the PM.

### Alanine scanning mutagenesis of LLP2/3 junction

Results above with CT truncation constructs suggested that the LLP2/3 region is required for Env enrichment in the condensed ERC, and also plays a role in regulating particle incorporation. We next sought to identify key residues or specific motifs that determine ERC localization and particle incorporation. We therefore performed alanine scanning mutagenesis within the C-terminus of LLP2 and N-terminal portion of LLP3 (throughout the region spanning L774 to N809, depicted in Figure 1C). Amino acids in this region of codon-optimized JRFL Env were mutated in pairs to alanines to ablate their functional groups, while existing alanine residues in this region were left unchanged. Each mutant was transfected into HeLa cells together with FIP1C_560-649_, and colocalization measured as before. Representative images are shown in Figure 4A. Quantitation of colocalization was performed from 20 images for each mutant, with the results shown in Figure 4B. The majority of alanine substitution introduced into the LLP2/3 junction had no or minimal effects on colocalization with FIP1C_560-649_ (represented by KY_794_AA, Figure 4A). However, five mutants demonstrated significantly reduced colocalization, as shown in Figure 4B, with representative images in Figure 4A. From these, we selected the three most N-terminal mutants for evaluation in the context of provirus, in order to further elucidate the role of the LLP2/LLP3 junction. These residues are depicted in Fig. 1D and E, derived from the published CT baseplate structure (19). These included a dileucine motif (LL_784_) at the C-terminus of LLP2, a highly conserved tryptophan glutamate dyad (WE_790_) motif at the N-terminus of LLP3, and a dual tryptophan (WW_796_) seven amino acids into LLP3 (overlapping with the previously described YW_795_ motif implicated in FIP1C-mediated Env trafficking (23)).

**FIG 4.**
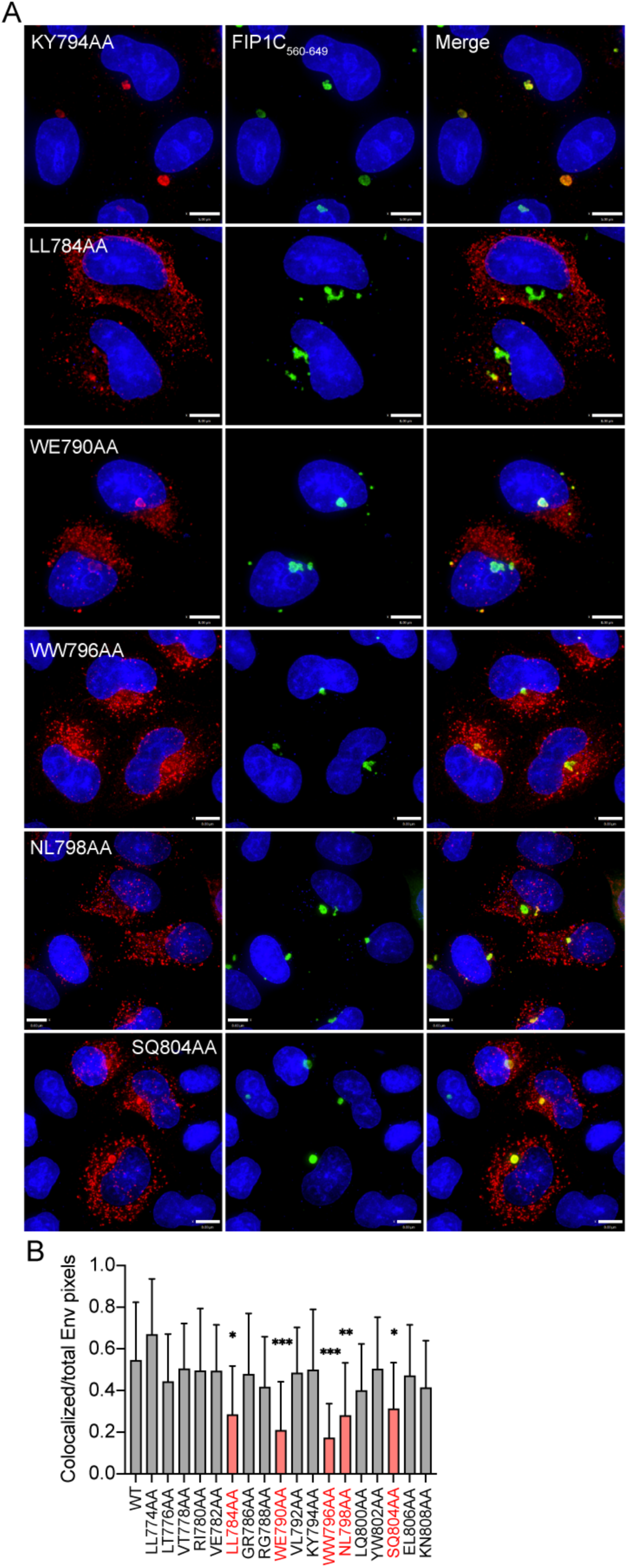
Mutations in LLP2 and LLP3 substantially reduce colocalization with FIP1C_560-649_. (A) Colocalization of Env point mutants with FIP1C_560-649_. KY794AA represents a mutant with WT degree of colocalization. All other mutants shown are significantly less colocalized with FIP1C_560-649_. Bar = 8 μm. (B) Colocalization of WT or mutant Env with GFP-FIP1C_560-649_ was quantified from 20 cells per Env construct; Manders colocalization coefficient is reported as mean ± SD. Mutants with significantly reduced colocalization are highlighted in red. Significant differences vs WT were assessed using one-way ANOVA with Dunnett correction for multiple comparisons. *, P<0.05; **, P<0.01; ***, P<0.001.

### Effect of LLP2/3 mutations in context of NL4-3 virus

The three dual-residue motifs identified above were next mutated within the context of NL4-3 provirus individually and in combination. In order to maintain amino acid identity within the *rev* open reading frame, we were unable to employ alanine codons for two of these motifs but instead generated substitutions to residues other than alanine (LL_784_RR, and WW_796_LS), while the WE_790_AA substitution was introduced as in the Env expression studies above. While assessing colocalization of NL4-3 Env mutants with the ERC, we observed that the condensed ERC generated upon GFP-FIP1C_560-649_ expression was frequently disrupted in transfected cells, resulting in a more diffuse cytoplasmic distribution of the GFP signal (compare 5A and 5B). This was true even in the absence of Env (Figure 5B, NL ΔEnv, right images), suggesting that another viral factor was causing the loss of condensation of the recycling compartment. Quantification of the area of diffuse GFP-FIP1C_560-649_ signal from infected cells vs FIP1C_560-649_ alone demonstrated the degree to which recycling membranes are disrupted during proviral expression (Figure 5C). Approximately 10% of infected cells displayed highly condensed FIP1C_560-649_ compartments (Figure 5D), and we chose to assess Env colocalization with FIP1C_560-649_ only in those cells.

**FIG 5.**
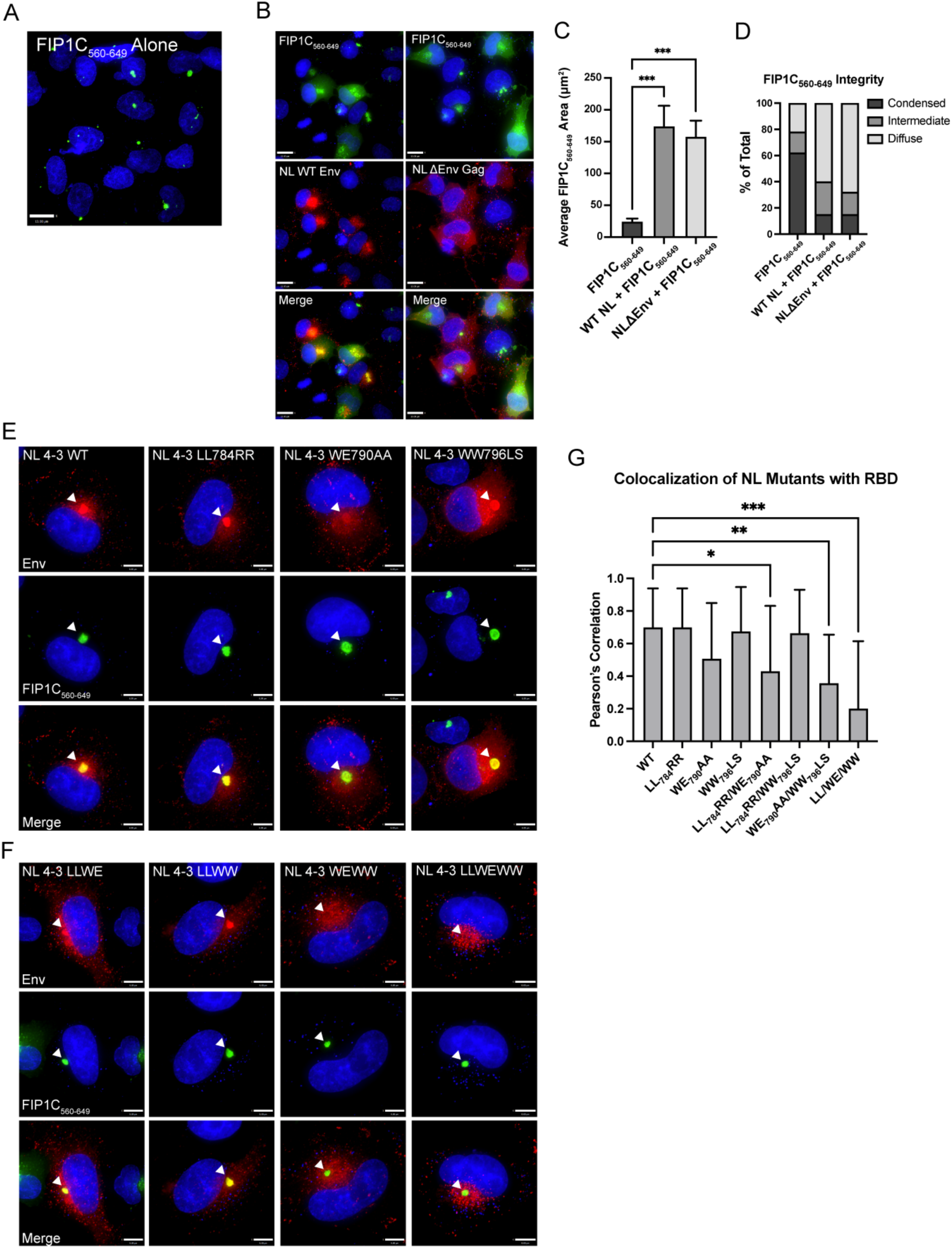
NL4-3 disrupts the condensed ERC; evaluation of mutations that disrupt Env colocalization with the ERC in proviral context. (A) Distribution of GFP-FIP1C_560-649_ alone. (B) Distribution of FIP1C_560-649_ with either NL4-3 WT or NL4-3 ΔEnv. Cells were fixed at 48 hours post transfection, stained for Env (NL WT) or for Gag (NL4-3 ΔEnv) and imaged. (C) GFP-FIP1C_560-649_ expressing cells from A, or infected HeLa cells from B were defined as a region of interest and the area of GFP-FIP1C_560-649_ was quantified within those ROIs. Results of area calculations are reported from 50 cells. (D) Cells from (C) were ranked based on the shape of the GFP-FIP1C_560-649_ compartment, with results reported as either condensed, intermediate, or diffuse. (E) Representative images from colocalization studies of individual point mutants in HeLa cells coexpressing NL4-3 provirus and GFP-FIP1C_560-649_. (F) Distribution of NL4-3 Env bearing two combined mutations or all three point mutations from LLP2/LLP3 junction in cells coexpressing GFP-FIP1C_560-649_. Arrowheads emphasize Env in the ERC. (G) Quantification of colocalization of NL4-3 Env with GFP-FIP1C_560-649_. Results are reported as mean ± SD of Pearson’s correlation coefficient for 20 cells per condition. Results were analyzed by one-way ANOVA with Dunnett correction. *, P<0.05; **, P<0.01; ***, P<0.001.

Individually, substitutions that had been found to be defective in ERC localization when expressed as Env alone had less of an effect in reducing ERC localization when expressed in intact virus (Figure 5E and 5G). To further examine the role of the residues surrounding the LLP2/LLP3 junction in Env incorporation, we generated combinations of two (LL_784_RR/WE_790_AA; LL_784_RR/WW_796_LS; and WE_790_AA/WW_796_LS) or all three (LL_784_RR/WE_790_AA/WW_796_LS) paired amino acid substitutions. LL_784_RR/WW_796_LS maintained ERC localization, not significantly different than the individual paired substitutions. Remarkably, however, WE_790_AA/WW_796_LS and LL_784_RR/WE_790_AA/WW_796_LS displayed significantly reduced ERC concentration as measured by colocalization with condensed FIP1C_560-649_ in 20 cells (Figure 5F and 5G). Additionally, LL_784_RR/WE_790_AA also appeared to be weakly reduced relative to WT (Figure 5G). Thus we conclude that the WE_790_AA/WW_796_LS mutant in the viral context reproduced the phenotype originally mapped by loss of ERC localization through truncation and deletion constructs, suggesting an important contribution for these tryptophan-based motifs within LLP3 in ERC localization.

Continuing the logic employed in the truncation studies above, we hypothesized that the dual tryptophan pair mutation that disrupted ERC localization (WE_790_AA/WW_796_LS) would result in diminished Env incorporation into particles. When particles were produced from Hela cells, single motif substitutions LL_784_RR and WE_790_AA had modest effects on Env particle incorporation as measured by gp41/p24 ratio, while WW_796_LS was reduced to 55% of WT (Figure 6A and 6C). The dual paired mutants LL_784_RR/WE_790_AA and LL_784_RR/WW_796_LS were incorporated to nearly the same extent as the WW_796_LS mutation (56% and 45% of WT respectively). However, combining substitutions of both tryptophan containing motifs (WE_790_AA/WW_796_LS) resulted in a much more robust reduction in Env incorporation at 32% of WT. The triple mutant, LL_784_RR/WE_790_AA/WW_796_LS, demonstrated reduced Env incorporation similar to WE_790_ AA/WW_796_LS (Figure 6A and 6C), suggesting that the tryptophan motifs were the critical determinants for particle incorporation and not the dileucine motif. The level of Env particle incorporation for the dual tryptophan motif mutant and for the triple motif mutant was reduced more substantially when particles were harvested from infected H9 T cells (Figure 6B and 6D). These results support the idea that mutations in LLP3 that disrupt ERC localization result in a corresponding reduction in particle incorporation of Env.

**FIG 6.**
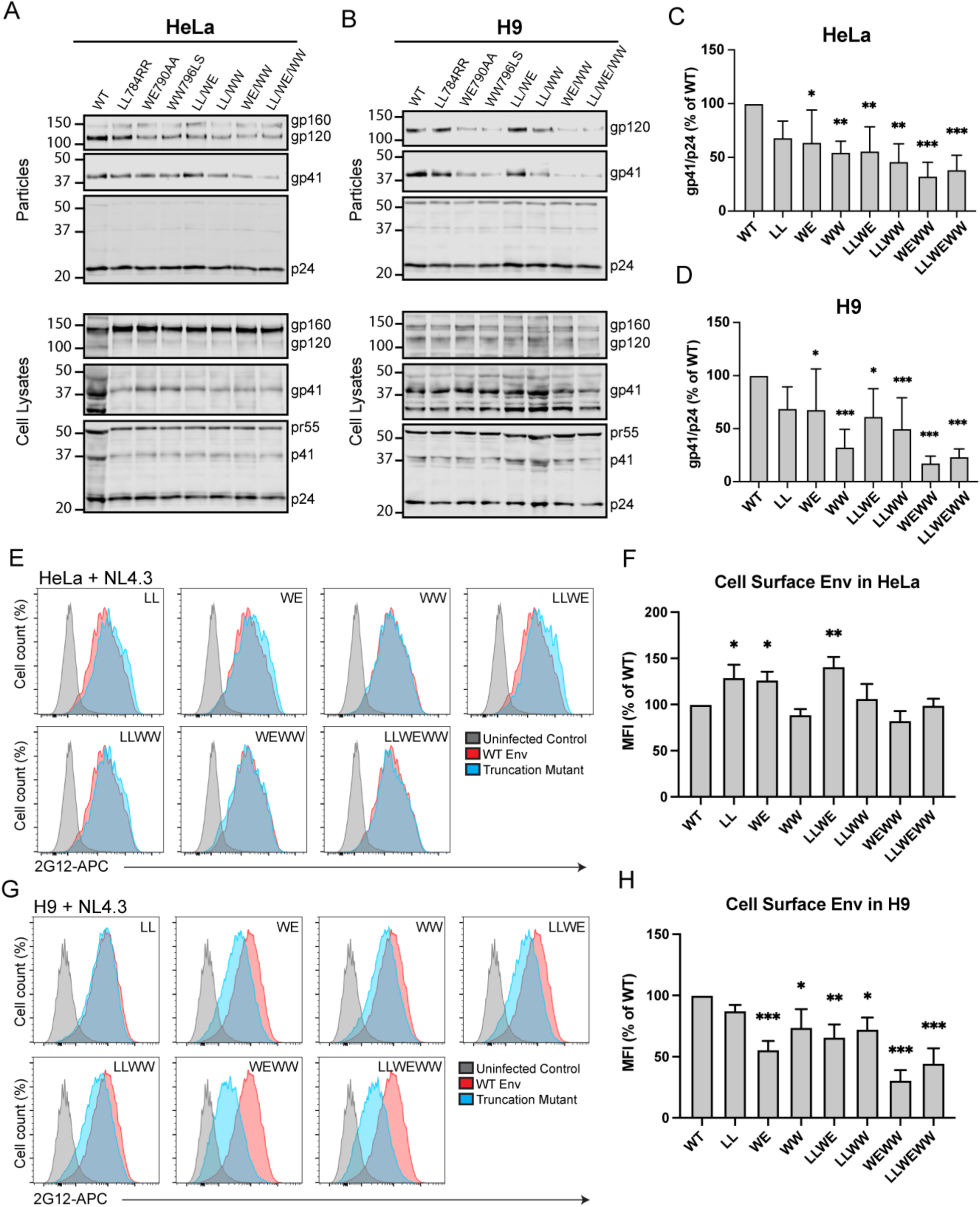
Tryptophan motifs in LLP3 mediating Env ERC localization are crucial for particle incorporation. (A) Transfected HeLa cells and (B) H9 cells infected with VSV-G-pseudotyped NL4-3 virus encoding mutant Envs were used to measure Env incorporation. Viral supernatants were pelleted through a sucrose cushion, normalized for virus production by p24 ELISA, and blotted. Westerns were analyzed by background subtracted densitometry and gp41/p24 ratios from four experiments for HeLa and seven experiments for H9 are reported as percent of WT (C and D). (E) Transfected HeLa cells expressing NL4-3 with the indicated Env point mutant were analyzed for cell surface Env content using flow cytometry. Cells were fixed and stained with 2G12 anti-gp120 directly conjugated to APC followed by permeabilization and staining for Gag. Gag positive cells were analyzed for cell surface Env. (F) Flow data in E is reported as mean ± SD of MFI as % of WT from three experiments. Significance was assessed using one-way ANOVA with Dunnett correction for multiple comparisons. (G) H9 cells were infected with pseudotyped NL4-3 containing the indicated point mutation, then were processed as described in E. H) Flow data in G is reported as described in F. *, P<0.05; **, P<0.01; ***, P<0.001.

Disruption of CT motifs involved in recycling might be expected to reduce cell surface levels of Env. However, in transfected HeLa cells a significant reduction was not observed, but instead there was a modest increase in cell surface Env for three mutants (LL_784_RR, WE_790_AA and LL_784_RR/WE_790_AA, Figure 6E and 6F). The dual tryptophan motif substitution WE_790_AA/WW_796_LS was reduced slightly but this reduction did not reach statistical significance (Figure 6F). In H9 cells the trend was substantially different, as each of the panel of mutants except LL_784_RR were significantly reduced on the cell surface of infected cells. WE_790_AA/WW_796_LS and LL_784_RR/WE_790_AA/WW_796_LS showed the most robust reduction, with only 26% and 32% of WT cell surface Env, respectively (Figure 6G and 6H). Thus, in H9 cells a significant reduction in cell surface Env was observed for those mutants that failed to localize with the ERC, perhaps a reflection of diminished recycling to the PM.

We anticipated that the LLP2/3 junction mutants that incorporated Env poorly would demonstrate defects in particle infectivity and in multiple-round replication assays. For the individual mutations LL_784_RR and WE_790_AA, and for the two dual motif mutants, LL_784_RR/WE_790_AA, and LL_784_RR/WW_796_LS, we observed a reduction in infectivity corresponding with the loss of incorporation described previously (Figure 7A). In agreement with Env incorporation studies above, we observed a more significant and robust reduction in infectivity for WE_790_AA/WW_796_LS and LL_784_RR/WE_790_AA/WW_796_LS viruses (Figure 7A). We then examined the LLP2/3 mutants in multiple round/spreading infection assays in H9 cells. Viruses containing the single motif substitution LL_784_RR, WE_790_AA and WW_796_LS demonstrated significantly impaired replication (Figure 7B). Notably, combining these substitutions in WE_790_AA/WW_796_LS and LL_784_RR/WE_790_AA/WW_796_LS resulted in a more severe defect in replication in H9 cells (Figure 7C). LL_784_RR/WE_790_AA and LL_784_RR/WW_796_LS showed an intermediate degree of delay of replication as compared with the viruses containing substitutions in both tryptophan-based motifs. Together, these results implicate two tryptophan-based motifs located at the very N-terminus of LLP3, WW_796-797_ and WE_790-791_, in ERC localization, Env incorporation, and viral replication.

**FIG 7.**
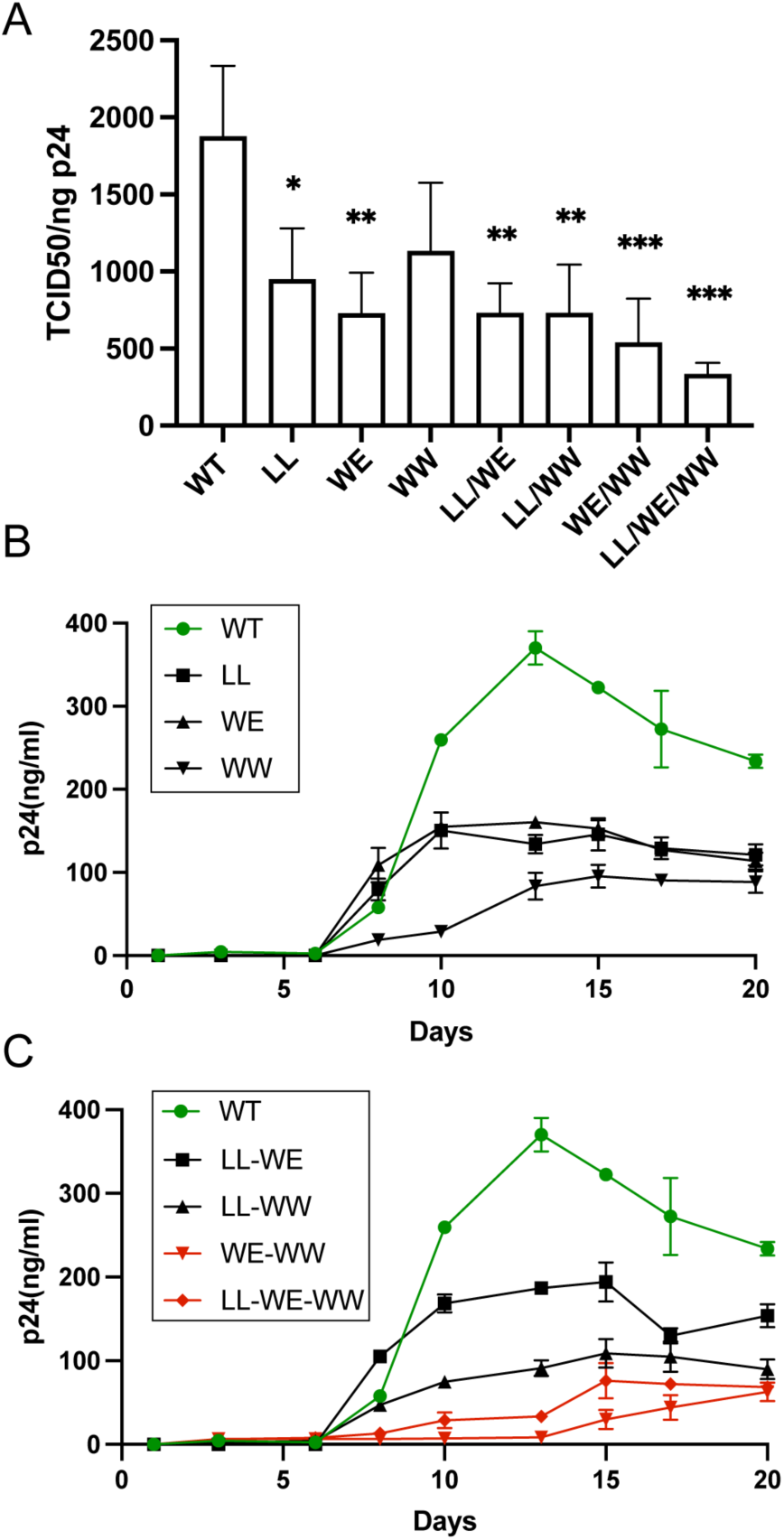
Trafficking-defective Env viruses are poorly infectious and exhibit diminished replication in spreading infection assays. (A) Tzm-bl reporter cells were incubated with viruses produced from H9 cells infected with WT or mutant viruses. After 48h, cells were lysed with the Britelite luciferase reagent and were analyzed by luminescence. TCID50 was normalized to p24 levels in viral supernatants as determined by p24 ELISA. (B) H9 cells were infected with NL4-3 bearing WT or mutant Envs indicated in the figure. Viral replication kinetics were assessed by monitoring p24 levels in supernatants over three weeks. (C) Combinations of the three substitution mutants were created in NL4-3 virus. H9 cells were infected as in (B). Red symbols/curves denote the mutants containing dual tryptophan mutations in the N-terminus of LLP3.

## DISCUSSION

The HIV-1 envelope glycoprotein takes an unusual path to reach the PM site of assembly. After delivery to the PM, Env trimers are rapidly endocytosed, entering the early/sorting endosome (5, 6, 32). From this location, membrane proteins can be sorted via rapid or slow recycling pathways back to the plasma membrane, or can be retained in maturing endosomes that eventually fuse with the lysosome. An alternative pathway mediated by the retromer complex can direct retrograde transport of Env to the Golgi (13). The pathways taken by HIV-1 Env from the early/sorting endosome remain incompletely defined. We have described a prominent role for the ERC in mediating Env incorporation, a pathway that was elucidated through study of the trafficking adaptor FIP1C (10, 11). These studies suggested a model in which directed recycling from the ERC, mediated by motifs within an intact CT and involving specific host adaptors, is required for Env incorporation into developing particles. An intriguing aspect of this line of investigation was the identification of a mutant in the Env CT that was deficient in Env incorporation and replication, YW_795_SL (23). This mutation within the proximal portion of LLP3 rendered virus deficient in incorporation in nonpermissive cell types, while not affecting incorporation in permissive cells. This suggested that this region of the CT is critical for cell type-specific particle incorporation, and that it may be involved in the recycling of Env to the PM site of assembly. Other studies have also pointed to a key role of the N-terminal portion of LLP3 in Env incorporation (21, 22). However, a model in which directed trafficking or co-targeting of Env to the particle assembly site is required for particle incorporation is one of several potential models for Env incorporation as proposed by the Freed laboratory (2). Another leading model to explain the requirement for the CT is the direct interaction model. A direct interaction between the CT and MA has been supported by several lines of evidence. Two biochemical studies have reported direct binding between MA and CT (33, 34). Suggestive evidence for direct Gag-Env CT interaction comes from the fact that maturation is required in order to “liberate” the CT from Gag and allow viral fusion, and that this constraint on fusion posed by the uncleaved Gag lattice can be relieved by truncation of the CT (35–39).

The goal of the present study was to identify specific motifs in the Env CT required for efficient recycling and particle incorporation, using ERC localization as the phenotype to identify recycling-competent Env. Using a truncated form of FIP1C as a probe for ERC localization, we found that truncation of the CT within LLP2, resulting in loss of the C-terminal portion of LLP2 and all of LLP3 and LLP1 (ΔCT82), resulted in complete loss of ERC localization. This truncation mutant links the findings of ERC localization with incorporation into particles, as the loss of trafficking to the ERC was associated with loss of particle association, despite Env being present at the cell surface at levels comparable to WT. The importance of the LLP2/3 junction was further highlighted by findings with a small internal deletion spanning this junction (Δ786-801), a deletion that resulted in Env exclusion from the ERC. Alanine-scanning mutagenesis then identified leucine-and tryptophan-based motifs at the LLP2/LLP3 junction that were important for ERC localization when Env and FIP1C_560-649_ were coexpressed. Upon introduction into intact provirus, the effects of single motif disruption were incomplete, while disruption of two tryptophan-based motifs, WE_790-791_ and WW_796-797_, resulted in a significant loss of Env incorporation, particle infectivity, and reduction in replication in multiple round replication assays. Together, these results strongly suggest a connection between CT-dependent Env localization in the ERC and incorporation into HIV-1 particles.

This study corroborates the importance of the N-terminal segment of the LLP3 region to Env particle incorporation that has been described in a number of previous studies (20, 21, 31, 40). A novel aspect of this study is that it demonstrates that ERC localization as determined in HeLa cells and particle incorporation in nonpermissive cell lines such as H9 are linked. The finding that ERC trafficking correlates with particle incorporation provides more support for a model in which CT-dependent recycling of Env trimers directed by host cell-specific factors is required for particle incorporation. This would be most consistent with a co-targeting model of Env incorporation, in which Gag and Env are delivered to the site of particle assembly on the PM via different routes (2). Notably, the co-targeting/recycling model remains compatible with a requirement for CT-MA interactions, as Env delivery to the site of Gag particle assembly may be a prerequisite to direct interactions.

We noted here as others have that partial Env CT truncations in HeLa cells which preserved LLP2 (represented here by ΔCT71 and ΔCT82) inhibited Env incorporation into particles, while deletion of all or a portion of LLP2 allowed more efficient Env incorporation. A similar phenotype had previously been described by the Aiken lab with their truncation studies (31). This finding was unique to this semipermissive cell line, as incorporation of Env with all of the truncations studied here was significantly reduced in H9 cells. We propose that this can be explained by competing pathways of intracellular trafficking, with the N-terminal portion of LLP3 involved in Env recycling to the plasma membrane, while LLP2 includes one of two motifs known to direct Env trafficking to the TGN in a retromer-dependent manner (13). Thus, truncation eliminating LLP3 while preserving LLP2 shunts Env away from assembly sites and into internal compartments, while truncations that remove all of LLP2 and LLP3 result in an increased quantity of Env on the plasma membrane, which may enhance Env incorporation through a passive incorporation mechanism (2). In nonpermissive cells such as H9, this passive or secondary incorporation mechanism is not present, so that enhanced cell surface envelope concentration results in no increase in Env incorporation into particles. Although this interpretation is admittedly likely to be an oversimplification, it provides a working model that ties trafficking mediated by LLP2 or LLP3 to the observed differences in cell type-specific Env incorporation.

It will be important to further define the recycling function mediated by the identified tryptophan-based motifs within the N-terminal portion of LLP3. Note that the WW_796_ substitutions studied here include alteration of the same tryptophan residue identified in the YW_795_SL mutant as important for Env incorporation and replication in nonpermissive cell lines (23). Both of these studies identified this key portion of the CT through screening mutants using assays related to intracellular trafficking; the first through a loss of cytoplasmic relocalization of FIP1C by YW_795_SL Env (23), and the present study through a completely different screening strategy employing ERC colocalization of Env. It is not clear at this time if the tryptophan-based motifs contribute to Env trafficking through direct interactions with host cell trafficking molecules or through a more indirect role. However, this study emphasizes that this small region of LLP3 is critical for Env incorporation into particles, and provides further support for a model in which the N-terminal portion of LLP3 is important for recycling of Env from the ERC to the site of particle assembly on the PM.

## MATERIALS AND METHODS

### Cells and plasmids

HeLa cells were obtained from ATCC (CCL-2). 293T cells were obtained from ATCC (CRL-3216). TZM-bl cells were obtained through the NIH HIV Reagent Program, Division of AIDS, NIAID, NIH (ARP-8129), contributed by Dr. John C. Kappes and Dr. Xiaoyun Wu. HeLa, 293T, and TZM-bl cells were grown in DMEM (Dulbecco’s Modified Eagle’s Medium) supplemented with 10% fetal bovine serum (FBS), 2 mM L-glutamine, 100 IU penicillin, and 100 μg/mL streptomycin. H9 T-lymphoid cells were obtained from ATCC (HTB-176) and were grown in RPMI (Roswell Park Memorial Institute) 1640 Medium supplemented with 10% FBS and 2 mM L-glutamine, 100 IU penicillin, and 100 μg/mL streptomycin. Strain NL4-3 Infectious Molecular Clone (pNL4-3), ARP-2852 was obtained through the NIH HIV Reagent Program, Division of AIDS, NIAID, NIH: Human Immunodeficiency Virus 1 (HIV-1), contributed by Dr. M. Martin. pNL4-3 CTΔ144 was provided by Eric Freed at NCI Viral Replication and Dynamics Program, Frederick, MD. GFP-FIP1C_560-649_ and codon-optimized JR-FL Env expression plasmids have been described previously (41).

### Cloning of Env deletion and substitution mutants

PCR was performed on Bio-Rad T100 Thermal Cycler with the primers outlined in Table S1 (supplemental materials). Amplified PCR products for truncation mutations in codon optimized Env were digested and ligated between EcoRI and NotI sites in pcDNA5/TO (Addgene). Ligated plasmid DNA was then transformed into JM109 competent cells (Promega). Site-directed mutagenesis of codon-optimized Env was performed using Quikchange (Agilent Technologies, Santa Clara, CA) followed by digestion with DpnI and transformation into JM109 cells. Truncation mutations in NL4-3 were created by subcloning the region of NL4-3 between EcoRI to XhoI into pBlueScript KS(-) (Addgene), followed by site-directed mutagenesis to introduce a stop codon to generate the desired truncation mutation, and then reinsertion into the NL4-3 backbone and transformation into Stbl4 electrocompetent cells (Thermo Fisher Scientific, Waltham, MA). Point mutations were inserted into NL4-3 from ordered gene fragments containing the desired mutations amplified by the amplification primers listed in Table 1 and ligated between the NheI to XhoI region of NL4-3, ligating these fragments into the NL4-3 backbone, and transforming into Stbl4 cells.

### Immunofluorescence microscopy

Cells were fixed for imaging with 4% paraformaldehyde in PBS for 15 minutes followed by three PBS washes. Permeabilization was performed using 0.2% Triton X-100 diluted in PBS for 5 minutes followed by three washes with PBS. Cells were blocked using Dako protein block (Agilent Technologies) for 30 minutes, washed once with PBS, then incubated with primary antibody diluted in Dako Antibody Diluent (Agilent Technologies) for one hour. Primary antibodies included 2G12 for Env, KC57 for Gag. Cells were washed thrice with PBS buffer containing 1% BSA and 0.05% IGEPAL CA-630 (Sigma Aldrich, St. Louis, MO), and then were stained with secondary antibody diluted in Dako Antibody Diluent. Secondary antibody for Env was anti-human Alexa Fluor 647 (Thermo Fisher Scientific). Cells were then washed with PBS once, stained with DAPI at 5 μg/mL, and washed twice more with PBS. Cells were then imaged using DeltaVision widefield deconvolution microscope (Leica Microsystems, Wetzlar, Germany).

### Measurement of particle infectivity

Tzm-bl cells were seeded at a density of 1.5×10^4^ cells per well of a 96 well plate and incubated overnight at 37°C. Viral supernatants were diluted 1:5 in quadruplicate in DMEM media containing 16 μg/mL DEAE dextran in the first well of a 96 well plate. Samples were serially diluted from 1:5 in column one to 1:48,828,125 in column 11. Diluted viral supernatants were then added to Tzm-bl cells and incubated at 37°C for 48 hours. Following incubation, 100uL of media was removed from the plate and incubated with 100 uL Britelite Plus reagent (PerkinElmer, Waltham, MA) for 2 minutes followed by mixing and transferring to a black 96 well assay plate. Samples were then analyzed using the Synergy Neo2 (BioTek Instruments, Winooski, VT).

### HIV spreading infection assay

Three million H9 cells were infected overnight with 150 ng of VSV-G-pseudotyped NL4-3 virus bearing WT or mutant Env. These cells were washed the following day and resuspended in one mL of RPMI 1640 media supplemented with 10% FBS and 2 mM L-glutamine, 100 IU penicillin, and 100 μg/mL streptomycin. Every two to three days 100 μL of media was removed to measure p24 production and replaced with 200 μL of fresh media. P24 in supernatants was quantified by antigen capture ELISA.

### Western blotting for viral proteins

Viruses were pelleted through a 20% sucrose cushion and resuspended in RIPA buffer, and cells were lysed with RIPA buffer. Both viral supernatants and cell lysates were normalized for p24 by ELISA prior to loading onto gel. 4-12% gradient Bis-Tris gels were purchased from Thermo Fisher Scientific or cast at time of use from 10% polyacrylamide in 0.36 M BisTris buffer pH 6.4. Env was detected with 2F5 human anti-gp41 (Polymun Scientific, Klosterneuburg, Austria) or 2G12 for gp120/gp160, both diluted 1:1000 in Intercept blocking buffer (Licor Biosciences, Lincoln, NE) with 0.15% Tween-20. Blots were normalized for p24 as measured by ELISA; Gag was blotted with mouse anti-p24 antibody CA183, also diluted 1:1000 in Intercept buffer with 0.15% Tween-20. Secondary staining was performed with Licor secondary antibodies diluted 1:5000 in the same blocking buffer solution.

### Flow Cytometry

For cell surface staining, HeLa cells were plated at a density of 400,000 per well of a six well plate and transfected with 400 ng of mutant Env. 24 hours post transfection, cells were stained with zombie violet dye (BioLegend/Perkin Elmer) diluted 1:500 in PBS, then fixed with 4% paraformaldehyde. Cells were washed twice with PBS then blocked with Dako blocking reagent, followed by staining on the cell surface with 2G12 anti-gp120 directly conjugated to APC (Abcam, Cambridge, UK; ab201807) diluted 1:100 in Dako antibody diluent for 2 hours. 2G12 was washed out with PBS twice followed by permeabilization with 0.2% Triton X-100 and staining for intracellular Gag with KC57-FITC (Beckman Coulter, Pasadena, CA; 6604665) diluted 1:100 in Dako antibody diluent. Cells were washed with PBS and resuspended in MACS buffer (Miltenyi Biotec, North Rhine-Westphalia, Germany) followed by analysis using BD FACS Canto II. Alternatively, 5 million H9 cells were infected overnight with appropriate viral mutant with 1ug of p24 in 1 mL to ensure a high degree of infection. H9 cells were washed once with PBS the next day and incubated with 2 mL RPMI for an additional 24 hours. The next day, cells were processed as described previously.

## ACKNOWLEDGMENTS

This work was supported by R01 AI150486.

The funders had no role in study design, data collection and interpretation, or the decision to submit the work for publication.

Flow cytometry was performed using the Cincinnati Children’s Hospital Medical Center (CCHMC) Flow Cytometry Core. DNA sequencing was performed at the DNA Sequencing core at CCHMC.

## REFERENCES

1. Bernstein HB, Tucker SP, Hunter E, Schutzbach JS, Compans RW. 1994. Human immunodeficiency virus type 1 envelope glycoprotein is modified by O-linked oligosaccharides. Journal of Virology 68:463–468. doi:10.1128/jvi.68.1.463-468.1994

2. Checkley MA, Luttge BG, Freed EO. 2011. HIV-1 Envelope Glycoprotein Biosynthesis, Trafficking, and Incorporation. Journal of Molecular Biology 410:582–608. https://doi.org/10.1016/j.jmb.2011.04.042

3. Willey RL, Bonifacino JS, Potts BJ, Martin MA, Klausner RD. 1988. Biosynthesis, cleavage, and degradation of the human immunodeficiency virus 1 envelope glycoprotein gp160. Proceedings of the National Academy of Sciences 85:9580–9584. doi:10.1073/pnas.85.24.9580

4. Hallenberger S, Bosch V, Angliker H, Shaw E, Klenk HD, Garten W. 1992. Inhibition of furin-mediated cleavage activation of HIV-1 glycoprotein gp160. Nature 360:358–61. 10.1038/360358a0

5. Rowell JF, Stanhope PE, Siliciano RF. 1995. Endocytosis of endogenously synthesized HIV-1 envelope protein. Mechanism and role in processing for association with class II MHC. The Journal of Immunology 155:473–488.

6. Ohno H, Aguilar RC, Fournier M-C, Hennecke S, Cosson P, Bonifacino JS. 1997. Interaction of Endocytic Signals from the HIV-1 Envelope Glycoprotein Complex with Members of the Adaptor Medium Chain Family. Virology 238:305–315. https://doi.org/10.1006/viro.1997.8839

7. Boge M, Wyss S, Bonifacino JS, Thali M. 1998. A Membrane-proximal Tyrosine-based Signal Mediates Internalization of the HIV-1 Envelope Glycoprotein via Interaction with the AP-2 Clathrin Adaptor *. Journal of Biological Chemistry 273:15773–15778. 10.1074/jbc.273.25.15773

8. Wyss S, Berlioz-Torrent C, Boge M, Blot G, Höning S, Benarous R, Thali M. 2001. The highly conserved C-terminal dileucine motif in the cytosolic domain of the human immunodeficiency virus type 1 envelope glycoprotein is critical for its association with the AP-1 clathrin adaptor [correction of adapter]. J Virol 75:2982–92. PMC115924.10.1128/jvi.75.6.2982-2992.2001

9. Byland R, Vance PJ, Hoxie JA, Marsh M. 2007. A conserved dileucine motif mediates clathrin and AP-2-dependent endocytosis of the HIV-1 envelope protein. Mol Biol Cell 18:414–25. PMC1783771.10.1091/mbc.e06-06-0535

10. Kirschman J, Qi M, Ding L, Hammonds J, Dienger-Stambaugh K, Wang J-J, Lapierre LA, Goldenring JR, Spearman P, Kirchhoff F. 2018. HIV-1 Envelope Glycoprotein Trafficking through the Endosomal Recycling Compartment Is Required for Particle Incorporation. Journal of Virology 92:e01893–17. doi:10.1128/JVI.01893-17

11. Qi M, Williams JA, Chu H, Chen X, Wang J-J, Ding L, Akhirome E, Wen X, Lapierre LA, Goldenring JR, Spearman P. 2013. Rab11-FIP1C and Rab14 Direct Plasma Membrane Sorting and Particle Incorporation of the HIV-1 Envelope Glycoprotein Complex. PLOS Pathogens 9:e1003278. 10.1371/journal.ppat.1003278

12. Hoffman HK, Aguilar RS, Clark AR, Groves NS, Pezeshkian N, Bruns MM, van Engelenburg SB. 2022. Endocytosed HIV-1 Envelope Glycoprotein Traffics to Rab14(+) Late Endosomes and Lysosomes to Regulate Surface Levels in T-Cell Lines. J Virol 96:e0076722. PMC9327703.10.1128/jvi.00767-22

13. Groppelli E, Len AC, Granger LA, Jolly C. 2014. Retromer regulates HIV-1 envelope glycoprotein trafficking and incorporation into virions. PLoS pathogens 10:e1004518–e1004518. 10.1371/journal.ppat.1004518

14. Rowell JF, Ruff AL, Guarnieri FG, Staveley O, Carroll K, Lin X, Tang J, August JT, Siliciano RF. 1995. Lysosome-associated membrane protein-1-mediated targeting of the HIV-1 envelope protein to an endosomal/lysosomal compartment enhances its presentation to MHC class II-restricted T cells. The Journal of Immunology 155:1818.

15. Weaver N, Hammonds J, Ding L, Lerner G, Dienger-Stambaugh K, Spearman P. 2023. KIF16B mediates anterograde transport and modulates lysosomal degradation of the HIV-1 envelope glycoprotein **bioRxiv** doi: https://doi.org/10.1101/2023.03.01.530732 https://doi.org/10.1101/2023.03.01.530732

16. Hunter E, Swanstrom R. 1990. Retrovirus envelope glycoproteins. Curr Top Microbiol Immunol 157:187–253. 10.1007/978-3-642-75218-6_7

17. Murphy RE, Samal AB, Vlach J, Saad JS. 2017. Solution Structure and Membrane Interaction of the Cytoplasmic Tail of HIV-1 gp41 Protein. Structure (London, England: 1993) 25:1708–1718.e5. 10.1016/j.str.2017.09.010

18. Piai A, Fu Q, Cai Y, Ghantous F, Xiao T, Shaik MM, Peng H, Rits-Volloch S, Chen W, Seaman MS, Chen B, Chou JJ. 2020. Structural basis of transmembrane coupling of the HIV-1 envelope glycoprotein. Nature Communications 11:2317. 10.1038/s41467-020-16165-0

19. Piai A, Fu Q, Sharp AK, Bighi B, Brown AM, Chou JJ. 2021. NMR Model of the Entire Membrane-Interacting Region of the HIV-1 Fusion Protein and Its Perturbation of Membrane Morphology. Journal of the American Chemical Society 143:6609–6615. 10.1021/jacs.1c01762

20. Bültmann A, Muranyi W, Seed B, Haas J. 2001. Identification of two sequences in the cytoplasmic tail of the human immunodeficiency virus type 1 envelope glycoprotein that inhibit cell surface expression. Journal of virology 75:5263–5276. 10.1128/JVI.75.11.5263-5276.2001

21. Murakami T, Freed EO. 2000. Genetic Evidence for an Interaction between Human Immunodeficiency Virus Type 1 Matrix and &#x3b1;-Helix 2 of the gp41 Cytoplasmic Tail. Journal of Virology 74:3548–3554. doi:10.1128/JVI.74.8.3548-3554.2000

22. Bhakta SJ, Shang L, Prince JL, Claiborne DT, Hunter E. 2011. Mutagenesis of tyrosine and di-leucine motifs in the HIV-1 envelope cytoplasmic domain results in a loss of Env-mediated fusion and infectivity. Retrovirology 8:37–37. 10.1186/1742-4690-8-37

23. Qi M, Chu H, Chen X, Choi J, Wen X, Hammonds J, Ding L, Hunter E, Spearman P. 2015. A tyrosine-based motif in the HIV-1 envelope glycoprotein tail mediates cell-type&#x2013; and Rab11-FIP1C&#x2013;dependent incorporation into virions. Proceedings of the National Academy of Sciences 112:7575–7580. doi:10.1073/pnas.1504174112

24. Qi M, Chu H, Chen X, Choi J, Wen X, Hammonds J, Ding L, Hunter E, Spearman P. 2015. A tyrosine-based motif in the HIV-1 envelope glycoprotein tail mediates cell-type-and Rab11-FIP1C-dependent incorporation into virions. Proc Natl Acad Sci U S A 112:7575–80. PMC4475960.10.1073/pnas.1504174112

25. Blot G, Janvier K, Le Panse S, Benarous R, Berlioz-Torrent C. 2003. Targeting of the human immunodeficiency virus type 1 envelope to the trans-Golgi network through binding to TIP47 is required for env incorporation into virions and infectivity. J Virol 77:6931–45. PMC156179.10.1128/jvi.77.12.6931-6945.2003

26. Lambele M, Labrosse B, Roch E, Moreau A, Verrier B, Barin F, Roingeard P, Mammano F, Brand D. 2007. Impact of natural polymorphism within the gp41 cytoplasmic tail of human immunodeficiency virus type 1 on the intracellular distribution of envelope glycoproteins and viral assembly. J Virol 81:125–40. PMC1797254.10.1128/JVI.01659-06

27. Murakami T, Freed EO. 2000. The long cytoplasmic tail of gp41 is required in a cell type-dependent manner for HIV-1 envelope glycoprotein incorporation into virions. Proc Natl Acad Sci U S A 97:343–8. PMC26665.10.1073/pnas.97.1.343

28. Yu X, Yuan X, McLane MF, Lee TH, Essex M. 1993. Mutations in the cytoplasmic domain of human immunodeficiency virus type 1 transmembrane protein impair the incorporation of Env proteins into mature virions. Journal of Virology 67:213–221. doi:10.1128/jvi.67.1.213-221.1993

29. Akari H, Fukumori T, Adachi A. 2000. Cell-Dependent Requirement of Human Immunodeficiency Virus Type 1 gp41 Cytoplasmic Tail for Env Incorporation into Virions. Journal of Virology 74:4891–4893. doi:10.1128/jvi.74.10.4891-4893.2000

30. Anokhin B, Spearman P. 2022. Viral and Host Factors Regulating HIV-1 Envelope Protein Trafficking and Particle Incorporation. Viruses 14. PMC9415270.10.3390/v14081729

31. Jiang J, Aiken C. 2007. Maturation-Dependent Human Immunodeficiency Virus Type 1 Particle Fusion Requires a Carboxyl-Terminal Region of the gp41 Cytoplasmic Tail. Journal of Virology 81:9999–10008. doi:10.1128/JVI.00592-07

32. Egan MA, Carruth LM, Rowell JF, Yu X, Siliciano RF. 1996. Human immunodeficiency virus type 1 envelope protein endocytosis mediated by a highly conserved intrinsic internalization signal in the cytoplasmic domain of gp41 is suppressed in the presence of the Pr55gag precursor protein. Journal of Virology 70:6547–6556. doi:10.1128/jvi.70.10.6547-6556.1996

33. Cosson P. 1996. Direct interaction between the envelope and matrix proteins of HIV-1. EMBO J 15:5783–8. PMC452325.

34. Alfadhli A, Staubus AO, Tedbury PR, Novikova M, Freed EO, Barklis E. 2019. Analysis of HIV-1 Matrix-Envelope Cytoplasmic Tail Interactions. J Virol 93. PMC6803273.10.1128/JVI.01079-19

35. Jiang J, Aiken C. 2006. Maturation of the viral core enhances the fusion of HIV-1 particles with primary human T cells and monocyte-derived macrophages. Virology 346:460–8. 10.1016/j.virol.2005.11.008

36. Jiang J, Aiken C. 2007. Maturation-dependent human immunodeficiency virus type 1 particle fusion requires a carboxyl-terminal region of the gp41 cytoplasmic tail. J Virol 81:9999–10008. PMC2045384.10.1128/JVI.00592-07

37. Murakami T, Ablan S, Freed EO, Tanaka Y. 2004. Regulation of human immunodeficiency virus type 1 Env-mediated membrane fusion by viral protease activity. J Virol 78:1026–31. PMC368813.10.1128/jvi.78.2.1026-1031.2004

38. Wyma DJ, Jiang J, Shi J, Zhou J, Lineberger JE, Miller MD, Aiken C. 2004. Coupling of human immunodeficiency virus type 1 fusion to virion maturation: a novel role of the gp41 cytoplasmic tail. J Virol 78:3429–35. PMC371074.10.1128/jvi.78.7.3429-3435.2004

39. Wyma DJ, Kotov A, Aiken C. 2000. Evidence for a stable interaction of gp41 with Pr55(Gag) in immature human immunodeficiency virus type 1 particles. J Virol 74:9381–7. PMC112366.10.1128/jvi.74.20.9381-9387.2000

40. Alfadhli A, Staubus AO, Tedbury PR, Novikova M, Freed EO, Barklis E. 2019. Analysis of HIV-1 Matrix-Envelope Cytoplasmic Tail Interactions. Journal of Virology 93:e01079–19. doi:10.1128/JVI.01079-19

41. Kirschman J, Qi M, Ding L, Hammonds J, Dienger-Stambaugh K, Wang JJ, Lapierre LA, Goldenring JR, Spearman P. 2018. HIV-1 Envelope Glycoprotein Trafficking through the Endosomal Recycling Compartment Is Required for Particle Incorporation. J Virol 92:DOI: 10.1128/JVI.01893-17. PMC5809729.10.1128/JVI.01893-17

